# Heat*seq: an interactive web tool for high-throughput sequencing experiment comparison with public data

**DOI:** 10.1101/049254

**Authors:** Guillaume Devailly, Anna Mantsoki, Anagha Joshi

## Abstract

Better protocols and decreasing costs have made high-throughput sequencing experiments now accessible even to small experimental laboratories. However, comparing one or few experiments generated by an individual lab to the vast amount of relevant data freely available in the public domain might be limited due to lack of bioinformatics expertise. Though several tools, including genome browsers, allow such comparison at a single gene level, they do not provide a genome-wide view. We developed Heat*seq, a web-tool that allows genome scale comparison of high throughput experiments (ChIP-seq, RNA-seq and CAGE) provided by a user, to the data in the public domain. Heat*seq currently contains over 12,000 experiments across diverse tissue and cell types in human, mouse and drosophila. Heat*seq displays interactive correlation heatmaps, with an ability to dynamically subset datasets to contextualise user experiments. High quality figures and tables are produced and can be downloaded in multiple formats.

**Availability:** Web application: www.heatstarseq.roslin.ed.ac.uk/. Source code: https://github.com/gdevailly.

## 1 Introduction

High throughput sequencing is now becoming routine for many biological assays including transcriptome analysis through RNA sequencing (RNA-seq), or transcription factor (TF) binding sites identification through chromatin immuno-precipitation followed by sequencing (ChIP-seq). Additionally, collaborative projects such as Bgee (Bastian *et al.*,) ENCODE (Bernstein *et al.*, 2012), and Roadmap Epigenomics (Kundaje *et al.*, 2015) have generated genome-wide datasets across hundreds of cell types or tissues. Despite this large data being freely available in the public domain, the lack of computational tools accessible to experimental scientists with no or elementary computational skills prohibits the use of this data to its full potential for discovery.

Though genome browsers, including summary tracks provided by many consortia, are extremely useful to study a few genes, promoters, or single nucleotide polymorphisms, they lack the genome-wide overview. Only a few public resources such as the CODEX database (Sanchez-Castillo *et al.*, 2015a) and the BLUEPRINT GenomeStats tool (Zerbino *et al.*, 2014) allow a genome-wide comparison with the user data. We therefore developed Heat*seq, a free, open source, web application providing fast and interactive comparison against high throughput sequencing experiments in the public domain. Users can upload a processed text file containing either gene expression value (FPKM or TPM), peak coordinates, or peak coordinates and corresponding expression value (CPM) for CAGE. The application provides clustered correlation heatmaps, summarising global similarities between all samples in the dataset and the user sample. Heat*seq provides over 12,000 publicly available genome-wide experiments in human, mouse and drosophila for fast and interactive comparison. In summary, Heat*seq is an interactive web tool that allows users to contextualise their sequencing data with respect to vast amounts of public data in a few minutes without requiring any programming skills.

## 2 Methods

### 2.1 Data collection

We collected gene expression data (RNA-seq), TF ChIP-seq data and CAGE (Cap Analysis of Gene Expression) data (over 4000 individual experiments) from Bgee (Bastian *et al.*,) Blueprint epigenome (Pradel *et al.*, 2015), CODEX (Sánchez-Castillo *et al.*, 2015b), ENCODE (Bernstein *et al.*, 2012), FANTOM5 (Forrest *et al.*, 2014), FlyBase (Attrill *et al.*, 2016), GTEx (Lonsdale *et al.*, 2013), modENCODE (Celniker *et al.*, 2009) and Roadmap Epigenomics (Bernstein *et al.*, 2010), in human, mouse and drosophila (table 1). Data formatting was done using R (R scripts available on GitHub). Heatmaps represent Pearson's correlation values between experiments calculated using a Gene x Experiment numeric matrix with gene expression values for expression data (log scaled), a Genomic regions x Experiments binary matrix indicating presence or absence of a peak for TF ChIP-seq data and a Genomic regions x Experiments numeric matrix of expression values for CAGE data (log scaled). Importantly, we constructed a metadata table which provides a web-link to original data and allows users to sub select each dataset.

When the user uploads a file, we compute an approximation of correlation values between the file and every experiment in a dataset using the following rules:

- For HeatRNAseq, if a gene is present in the dataset but not in the user file, it is assigned a zero expression value. If a gene is present in the user file but not in the dataset, it will be ignored.
- For HeatChIPseq, if a peak in the user file does not overlap any regions in the dataset, it will be discarded.
- For HeatCAGEseq, if a CAGE peak in the user file does not overlap any CAGE peak in the dataset, it will be discarded.

**Table 1:** Datasets implemented in Heat*seq

| Assay | Dataset | Organism | Samples |
| --- | --- | --- | --- |
| RNA-seq | Bgee | human | 77 |
|  |  | mouse | 109 |
|  | Blueprint epigenome | human | 163 |
|  | ENCODE | human | 302 |
|  |  | mouse | 192 |
|  | Roadmap Epigenomics | human | 57 |
|  | GTEX (summary) | human | 53 |
|  | GTEX (all samples) | human | 8555 |
|  | Flybase | drosophila | 124 |
| TF ChIP-seq | ENCODE | human | 690 |
|  | CODEX | human | 238 |
|  |  | mouse | 651 |
|  | modENCODE | drosophila | 85 |
| CAGE | Fantom5 | human | 1058 |
|  |  | mouse | 490 |

### 2.2 Web-application development

Heat*seq is an R shiny open source interactive tool which computes correlation values between the user file and each experiment in a dataset. Detailed user instructions are on the application website.

## 3 Results

### 3.1 Application description

Heat*seq tool supports three data types: HeatRNAseq, HeatChIPseq and HeatCAGEseq. Data upload, correlation calculation and heatmap generation takes about a minute. Importantly, users can interactively sub select relevant experiments using the metadata information (e.g. cell type, TF name). The interactive heatmap also allows selecting different clustering methods as well as zooming in and out on the heatmap. The high resolution figures and tables can be downloaded in multiple formats. Thus, Heat*seq provides global overview of relationships between public experiments and the user data. Four user scenarios are discussed below.

**Figure 1:**
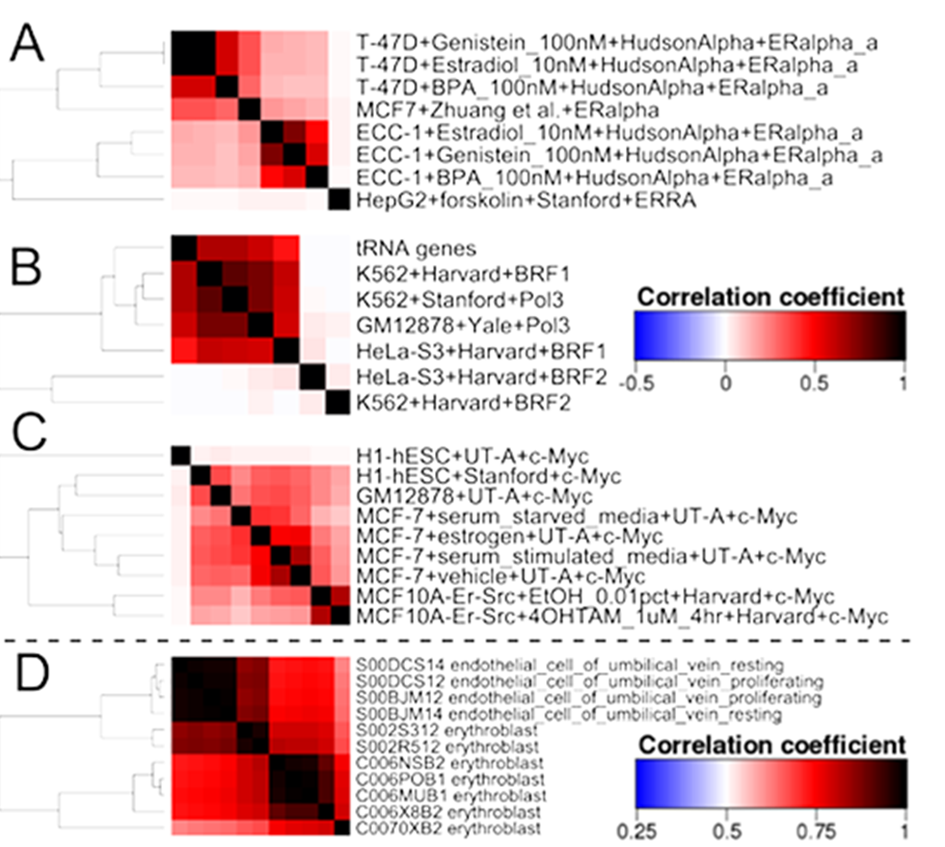
Correlations heatmaps from Heat*seq. **A.** ERα ChIP-seq in MCF7 cells from Zhuang et al. is closer to ENCODE ERa ChlP-seq in T-47D than in ECC-1 cells. **B.** BRF1 and RNA PolIII bind tRNA genes, but nor BRF2. **C.** c-MYC ChIP-seq in H1-hESC from UT-A and Stanford show low correlation. The color key next to B is for A, B and C. **D.** Two erythroblast RNA-seq samples from BLUEPRINT are closely related to endothelial cells.

### 3.2 User scenarios

#### 3.2.1 User data quality control

We compared a Neocortex, 10 days post-partum (Ray *et al.*, 2015) RNA-seq sample with Bgee mouse RNA-seq data using HeatRNAseq. The top five correlation values (PCC > 0.9) correspond to Bgee brain samples (Table 2). Thus, Heat*seq can be used as a fast data quality check for next generation sequencing data.

**Table 2:**
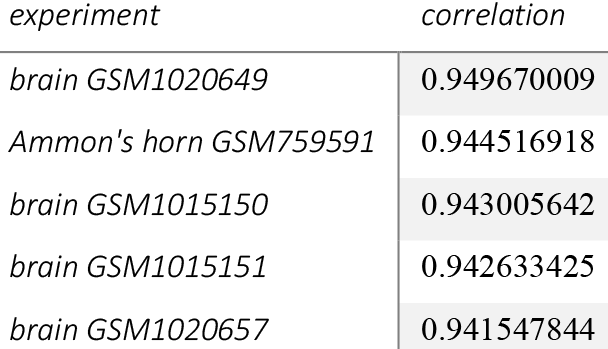
Top 5 experiments in the Bgee mouse RNA-seq dataset with the highest correlation value with a Neocortex RNA-seq experiment.

#### 3.2.2 Cell context identification

An oestrogen receptor (ER) alpha ChIP-seq in MCF7 cells (Zhuang *et al.*, 2015) comparison to the ENCODE TFBS dataset by sub-selecting ENCODE ER ChIP-seq experiments revealed that the binding pattern of ERα in MCF7 cells was more similar to its binding pattern in T-47D cells than in ECC-1 cells (Figure 1A). MCF7 and T-47D were derived from mammary tumours while ECC-1 is an endometrial cell line.

#### 3.2.3 New hypotheses by data integration

CpG islands (CGI) from the UCSC (Karolchik *et al.*, 2004) comparison to HeatChIPseq found that RNA polymerase II and TAF1 (Table 3) were enriched at CGIs, as about 50% of human gene promoters contain a CGI (Illingworth and Bird, 2009). Interestingly we identified factors avoiding CGIs including MAFK, GATA3 and ZNF274.

Similarly, tRNA promoters were highly correlated with RNA polymerase III, and its co-factors BDP1, RPC155, and BRF1 using HeatChIPseq. Interestingly, comparison with BRF family data revealed that BRF1, but not BRF2 was bound at tRNA genes (Figure 1B).

**Table 3:** Top and bottom 5 experiments in the ENCODE human TF ChIP-seq dataset with the highest and lowest correlation value with CpG island coordinates.

| experiment | correlation |
| --- | --- |
| A549+UT-A+Pol2 | 0.626574 |
| Gliobla+UT-A+Pol2 | 0.602679 |
| H1-hESC+HudsonAlpha+TAF1 | 0.602469 |
| ECC-1+DMSO_0.02pct+HudsonAlpha+Pol2 | 0.601939 |
| K562+Broad+PHF8_(A301-772A) | 0.600818 |
| ... | ... |
| A549+DEX_100nM+HudsonAlpha+FOXA1_(SC-101058) | -0.00415 |
| SH-SY5Y+USC+GATA3_(SC-269) | -0.00597 |
| MCF10A-Er-Src+4OHTAM_1uM_4hr+Harvard+c-Fos | -0.00613 |
| MCF-7+USC+GATA3_(SC-268) | -0.00924 |
| HepG2+Stanford+MafK_(ab50322) | -0.01513 |

#### 3.2.4 Public data assessment

Heat*seq can be used to assess data in the public domain, highlighted by two examples below amongst others:

A MYC ChIP-seq in H1-hESC cells does not cluster with other ENCODE MYC ChIP-seq experiments (Figure 1C), including H1-hESC sample from a different experimental group (Devailly *et al.*, 2015).

Two out of seven erythroblast RNA-seq samples from the Blueprint Epigenome consortium are more correlated with endothelial cells than with the rest of the erythroblast samples (Figure 1D).

## 4 Conclusion

With Heat*seq, comparing RNA-seq, ChIP-seq or CAGE experiments to hundreds of publicly available datasets becomes a trivial task. Researchers can now investigate the relationships between various high-throughput sequencing experiments fast and interactively without requiring any programming skills. Such analysis can assess data quality, cell variability, and generate novel regulatory hypotheses.

## Acknowledgements

We would like to thank Barry Horne for the R shiny server set-up and administration and the Edinburgh R user group (EdinbR) for their advice and support.

## References

Attrill, H. et al. (2016) FlyBase: establishing a Gene Group resource for Drosophila melanogaster. Nucleic Acids Res., 44, D786–D792.

Bastian, F. et al. Bgee: Integrating and Comparing Heterogeneous Transcriptome Data Among Species. In, Data Integration in the Life Sciences. Springer Berlin Heidelberg, Berlin, Heidelberg, pp. 124–131.

Bernstein, B.E. et al. (2012) An integrated encyclopedia of DNA elements in the human genome. Nature, 489, 57–74.

Bernstein, B.E. et al. (2010) The NIH Roadmap Epigenomics Mapping Consortium. Nat Biotech, 28, 1045–1048.

Celniker, S.E. et al. (2009) Unlocking the secrets of the genome. Nature, 459, 927–30.

Devailly, G. et al. (2015) Variable reproducibility in genome-scale public data: A case study using ENCODE ChIP sequencing resource. FEBS Lett.

Forrest, A.R.R. et al. (2014) A promoter-level mammalian expression atlas.Nature, 507, 462–70.

Illingworth, R.S. and Bird, A.P. (2009) CpG islands-’a rough guide'. FEBS Lett., 583, 1713–20.

Karolchik, D. et al. (2004) The UCSC Table Browser data retrieval tool. Nucleic Acids Res., 32, D493–6.

Kundaje, A. et al. (2015) Integrative analysis of 111 reference human epigenomes. Nature, 518, 317–330.

Lonsdale, J. et al. (2013) The Genotype-Tissue Expression (GTEx) project. Nat. Genet., 45, 580–5.

Pradel, L.C. et al. (2015) [The European Blueprint project: towards a full epigenome characterization of the immune system]. Médecine Sci. M/S, 31, 236–8.

Ray, S. et al. (2015) An Examination of Dynamic Gene Expression Changes in the Mouse Brain During Pregnancy and the Postpartum Period. G3 (Bethesda)., 6, 221–33.

Sánchez-Castillo, M. et al. (2015a) CODEX: a next-generation sequencing experiment database for the haematopoietic and embryonic stem cell communities. Nucleic Acids Res., 43, D1117–23.

Sánchez-Castillo, M. et al. (2015b) CODEX: a next-generation sequencing experiment database for the haematopoietic and embryonic stem cell communities. Nucleic Acids Res., 43, D1117–23.

Zerbino, D.R. et al. (2014) WiggleTools: parallel processing of large collections of genome-wide datasets for visualization and statistical analysis. Bioinformatics, 30, 1008–9.

Zhuang, T. et al. (2015) p21-activated kinase group II small compound inhibitor GNE-2861 perturbs estrogen receptor alpha signaling and restores tamoxifen-sensitivity in breast cancer cells. Oncotarget, 6, 43853–43868.

